# Identifying Effect Modification of Latent Population Characteristics on Risk Factors with a Sparse Varying Coefficient Regression

**DOI:** 10.1101/2024.11.30.626101

**Authors:** Ruofan Wang, Lei Fang, Yue Wang, Jin Jin

**Affiliations:** Department of Biostatistics, Epidemiology and Informatics, University of Pennsylvania, Philadelphia, PA 19104, USA; Department of Biostatistics and Data Science, University of Minnesota Twin Cities, Minneapolis, MN 55455, USA; Department of Biostatistics and Informatics, Colorado School of Public Health, 13001 E. 17th Place, Aurora, CO 80045, USA

**Keywords:** Effect modification, Latent features, Sparse penalized regression, Varying coefficient models

## Abstract

Leveraging observational data to understand the associations between risk factors and disease outcomes and conduct disease risk prediction is a common task in epidemiology. While traditional linear regression and other machine learning models have been extensively implemented for this task, the associations between risk factors and disease outcomes are typically deemed fixed. In many cases, however, such associations may vary by some underlying features of the individuals, which may involve certain subpopulation characteristics and environmental factors. While data for these latent features may not be available, the observed data on risk factors may have captured some proportion of the variation in these features. Thus extracting latent factors from risk factors and incorporating this effect modification into the model may better capture the underlying data structure and improve inference. We develop a novel regression model with some coefficients varying as functions of latent features extracted from the risk factors. We have demonstrated the superiority of our approach in various data settings via simulation studies. An application on a dataset for lung cancer patients from The Cancer Genome Atlas (TCGA) Program showed that our approach led to a 6% - 118% increase in (AUC-0.5) for distinguishing between different lung cancer stages compared to the classic lasso and elastic net regressions and identified interesting latent effect modifications associated with certain gene pathways.

## 1 Introduction

Classic parametric models, such as (generalized) linear regression and penalized regression models, are widely implemented in public health research to identify risk factors associated with various disease outcomes and predict risks of diseases. Linear regression models have been popular because of their model simplicity, well-established properties, interpretability, and robust performance on various data modalities. While advanced machine learning algorithms like deep neural networks (DNNs) are becoming increasingly popular for disease risk prediction, they typically require a sufficiently large number of samples to outperform classic regressions, and their black-box structures have posed long-standing challenges regarding algorithm interpretability and association analysis.

Nevertheless, classic regression models may have missed some important data information, where the associations between risk factors and the outcome may vary by some unknown factors or known factors with no available data. One example is the population heterogeneity in the genetic architectures of human traits and diseases. Recent work has shown extensive evidence of ancestral diversity in the effects of genetic variants on various diseases such as type 2 diabetes Smith et al. (2024), Alzheimer’s disease Lake et al. (2023), and cancers such as breast Hughes et al. (2022), prostate Uzamere et al. (2022), and testicular cancers Pyle and Nathanson (2016). Research has shown that accounting for ancestral heterogeneity in the effects of genetic variants by modeling ancestry-specific effects can lead to a fundamental improvement in causal variant detection and genetic risk prediction Kachuri et al. (2024); Zhang et al. (2023); Jin et al. (2024); Zhang et al. (2024). While clinical trials have linked microbiome with various diseases, individual microbiome studies often face replicability issues with inconsistent conclusions due to limited sample sizes, heterogeneous study samples, and varying data collection protocols Duvallet et al. (2017). Studies have associated microbial diversity with differences in lifestyles, such as diet and exercise, which may partially explain the inconsistent findings in microbiome-disease associations. Another example is the identification of disease-causal genes. While the causal genes may have universal effects on a disease outcome, they may function differently across different tissues and cell types, and there may be cofunctioning gene sets with cluster effects. Finally, all these problems suffer from the high dimensionality of the data, where the sample size is typically small compared to the number of risk factors. Most of the studies on these problems still implement regularized regressions assuming homogeneity among samples, which, however, lacks the power to detect potential effect heterogeneity and thus may lead to biased associations and underpowered predictions.

The varying coefficient model, which allows model coefficients to vary as smooth functions of other variables, has been studied to explore dynamic patterns of the exposure-outcome associations Hastie and Tibshirani (1993); Fan and Zhang (2008). The general varying coefficient model assumes a basic form of multivariate regression with the coefficients being unknown functions of some observed effect modifiers that need to be estimated. Varying coefficient models are flexible, allowing any form of the functions explaining the relationship between the effect modifiers and risk factors, and interpretable under a regression framework. There are also recent attempts to account for hidden confounding and effect modification in the high-dimensional contexts. For example, a high-dimensional mixed linear regression (MLR) was proposed to capture hidden effect heterogeneity by extracting binary latent group indicators from risk factors Zhang et al. (2020). Recent work also proposed a hidden confounding model, which assumes that there exist hidden confounders that are some hidden features of the risk factors and conduct inference by doubly debiased lasso Guo et al. (2022). The hidden confounding model, however, does not further consider effect modification of the hidden features. The setting with risk factors having varying-by-latent-factor associations has not been previously investigated. It is, however, common in reality, where the exposure-outcome associations can vary by some underlying features of the individuals, which are typically unknown or unavailable but may involve certain subpopulation characteristics and environmental factors that could affect the effect of the risk factors on disease risks.

In this study, we propose LF-VCR, a sparse varying coefficient regression to capture hidden effect modification by latent features extracted from the risk factors. We develop the model under a varying coefficient model framework but with latent effect modifiers instead of observed ones. The relationship between the latent features and the risk factors is hypothesized to be described by a general function *f* (*·*). For example, in microbiome studies, some latent features in the microbiome data are hypothesized to partially reflect lifestyle, such as diet and exercise, that may modify the effect of microbiome on individuals’ health status. In the absence of the observed data for these lifestyle characteristics, we may extract some latent factors from the observed data on risk factors to capture such hidden varying coefficient structures. With *f* being a linear function, the model basically adds interaction terms between the risk factors and their latent features. The latent features can be linear combinations of the risk factors generated by factor models or principal component analysis (PCA), or nonlinear features generated by deep-learning autoen-codersLi et al. (2023). We refer to the model corresponding to linear and nonlinear latent features by LF-VCR and LF-VCR (ae), respectively. Once we extract the latent factors, we then assume a sparse effect modification of the factors on the high-dimensional risk factors, i.e., a small proportion of the risk factors having coefficients that vary by the latent factors. We adopt a sparse-group lasso Simon et al. (2013) to model the effect modifications, assuming all latent factors simultaneously have either zero or nonzero effects on each risk factor. We show by simulation studies that the proposed LF-VCR and LF-VCR (ae) methods achieve higher predictive accuracy, lower estimation error, and better variable selection over classic penalized regressions like lasso and elastic net regressions. An application of our methods to model lung cancer stages by gene expression data utilizing data from The Cancer Genome Atlas (TCGA) Program shows that our method has led to an increase in AUC for distinguishing between lung cancer stage I and stage III from 0.581 (lasso) and 0.5992 (elastic net) to 0.6305, and an increase in AUC for distinguishing between lung cancer stage I and stage IV from 0.5388 (lasso) and 0.5418 (elastic net) to 0.5851. We have identified 9 common pathways that are highly correlated with the extracted latent factors. These pathways, linked to inflammation, immune modulation, and cellular signaling, play crucial roles in lung cancer biology by influencing key processes such as tumor initiation, growth, and immune evasion. Understanding these mechanisms can be critical for identifying potential therapeutic targets and developing strategies to predict and manage lung cancer progression.

## 2 Method

### 2.1 Notations and Model Setup

Let *y* denote the disease outcome of interest, **c** = (*c*_1_, …, *c*_*L*_)^*T*^ the *L*-dimensional covariates (e.g., age, gender), **x** = (*x*_1_, …, *x*_*p*_)^*T*^ the *p*-dimensional risk factors, and **z** = (*z*_1_, …, *z*_*q*_)^*T*^ the *q*-dimensional factors that summarize the lower-dimensional latent features embedded in the risk factors **x** that may capture variation in hidden variables, such as subpopulation structures and environmental factors that affect the associations between risk factors and outcomes. We denote the observed data of the variables *y*, **x, c** for a random sample of *N* individuals from a population *P* by 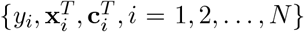. Since the latent features 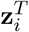 cannot be directly measured, we will utilize different approaches to estimate 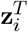.

Our goal is to develop a well-calibrated model *E*(*y*|**x, z, c**) within population *P*, describing variation in outcome *y* by the risk factors **x** and the latent factors **z** extracted from **x**. We assume, without loss of generality, that the outcome variable is continuous, but the method can be easily applied to disease outcomes in a generalized regression framework. Specifically, we propose to model *y*_*i*_ with **x, z**, and **c** by: 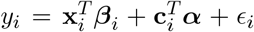, where 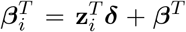, and **x**_*i*_ = Λ**z**_*i*_ + **u**_*i*_, *i* = 1, …, *N*. Here, **Λ** is the loading matrix estimated during factor analysis, ***β***^*T*^ and **δ** are the unknown parameters and matrix that we aim to estimate. We start by considering a linear latent feature scenario, where the observed risk factors **x**_*i*_ are modeled as a linear function of the latent features **z**_*i*_. In this case, **Λ** is obtained through high-dimensional factor analysis, representing the effect of **z**_*i*_ on **x**_*i*_. While **Λ** itself is not the primary focus, it serves as an intermediary for extracting **z**_*i*_. Alternatively, the latent feature **z**_*i*_ can also be nonlinear. In the nonlinear feature scenario, feature extraction may involve more flexible approaches such as autoencoders, where a specific structure for **Λ** does not exist. The corresponding exposure model under a nonlinear setting takes a different form, accommodating the complexity of the extracted features.

We will discuss how both linear and nonlinear latent features are generated in detail in Section 2.3. ***β*** denotes the constant effect of risk factors on disease outcomes that does not change with latent features and the matrix **δ** controls the magnitude of the influence of **z**_*i*_, **x**_*i*_ interaction on *y*_*i*_, and we denote **δ** by the interaction matrix. The varying effect of **x**_*ij*_ on *y*_*i*_ is adjusted by (**δ**_*j*1_, ···, **δ**_*jq*_ )^*T*^ ·(**z**_*i*1_, ···, **z**_*iq*_ ). Writing out our model in detailed form, we have:

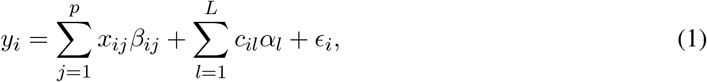

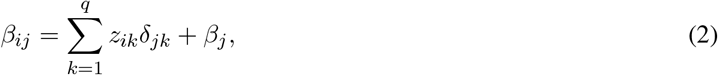

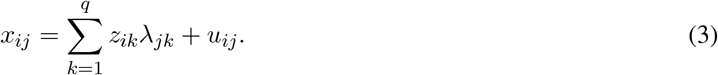

### 2.2 Objective Function with a Sparse-Group Lasso Penalty

Given the potential high dimensionality of the data, we induce sparsity on both the main effects of **x** and the interaction effects of **x** and **z** in model (1) with a sparse-group lasso penalty. This approach was chosen over traditional Lasso or Ridge penalties because it can better identify the risk factors that play the most significant roles in effect modifications. Using Lasso alone could result in all variables having non-zero estimates for at least one latent factor, leading to challenges in the interpretation of the varying coefficient patterns. By applying the sparse-group lasso penalty, we consider group structure by risk factor. This is, we assume the effect of each observed risk factor is either influenced by all latent features or by none of them. This means that **x**_*j*_**z**_1_, …, **x**_*j*_**z**_*q*_ will all have nonzero effects or all have no effect for each *j* = 1, …, *p*; in other words, **δ**_*j*1_, …, **δ**_*jq*_ are penalized as a group for each *j* = 1, …, *p*. For the main effects of the risk factors, (***β***_1_, ···, ***β***_*p*_), and covariate effects, (***α***_1_, ···, ***α***_*L*_), each of them is treated as an individual group and penalized separately. We thus have a total of *G* = 2*p* + *L* groups, with the first *p* groups corresponding to the interactions between **x**_*j*_s and **z**s, and the last *p* + *L* groups corresponding to the main effects of the *p* risk factors and *L* covariates. For simplicity of notations, we define 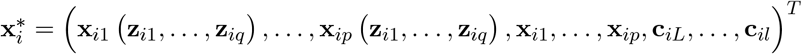 and let 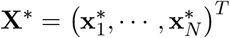 denote the row-wise concatenation of 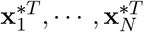. The corresponding vector ***β***^∗^ is then defined as: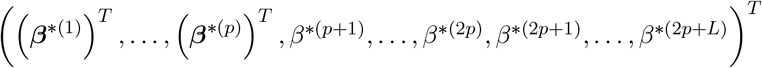. The correspondence of ***β***^∗^ with ***β*** and **δ** in (1) is that 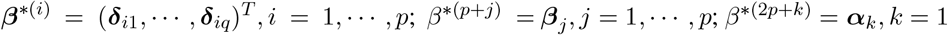.

With the sparse-group lasso penalty (Simon and Tibshirani, 2013), the objective function we want to minimize becomes:

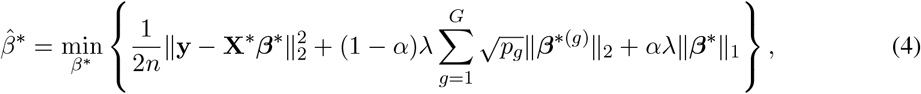

where ∥ ·∥_2_ denotes the Euclidean (ℓ_2_) norm, ∥ ·∥_1_ denotes the ℓ_1_ norm. Here, *α* and *λ* are two hyper-parameters. *α* ∈ [0, 1] controls the balance between the group lasso and the lasso penalties. Specifically, when *α* = 0, the penalty reduces to the group lasso, which enforces sparsity only at the group level and when *α* = 1, the penalty reduces to the standard lasso without considering group structure. *λ* determines the strength of regularization. A smaller *λ* will induce minimization of the squared error while a larger *λ* will emphasize variable selection.

### 2.3 Latent Feature Extraction

A major task in our method is to extract latent features **z** from the risk factors **x** to capture hidden population structures and environmental factors that may affect the exposure-outcome relationships. We propose two approaches to this task by extracting either linear or nonlinear latent features. The first approach utilizes a factor model assuming the observed risk factors are linear combinations of these latent factors, **x**_*i*_ = **Λz**_*i*_+**u**_*i*_, and obtains **z** through traditional statistical methods. The second approach does not assume any specific structure on the relationship between **x** and **z**, and applies a deep learning algorithm to extract feature representations. For the first approach, we propose feature extraction through Principal Orthogonal Complement Thresholding (POET) Fan et al. (2013), which is designed for sparse covariance estimation while extracting underlying factor structures. It models the observed exposure data as **x**_*i*_ = **Λz**_*i*_ + **u**_*i*_, where **Λ** is the factor loading matrix, **z**_*i*_ ∼ N (**0, I**_*K*_) represents latent factors, and **u**_*i*_ ∼ *N* (**0, Σ**_*u*_) denotes residual noise. It assumes the covariance matrix can be decomposed as **Σ** = **ΛΛ**^*T*^ + **Σ**_*u*_, where **ΛΛ**^*T*^ captures the low-rank structure and **Σ**_*u*_ represents sparse noise. POET uses a nonparametric estimator of **Σ** based on principal component analysis. Let 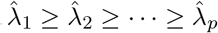 be the ordered eigenvalues of the sample covariance matrix 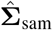 _sam_, and 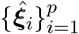 be their corresponding eigenvectors. Then the sample covariance has the following spectral decomposition: 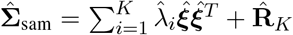 where 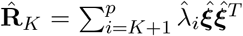 is the principal orthogonal complement. Therefore, 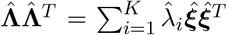. Inserting the estimated loading matrix, we could get the representation for latent factors 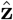

An autoencoder is a type of artificial neural networks (ANN) used to perform black-box feature extraction by mapping the input to latent representations and reconstructing the input, **X** → **Z** → **X**. These models extract nonlinear underlying features from the observed data and are widely used for unsupervised feature learning Hinton and Salakhutdinov (2006). For our second approach to extracting complex, structured latent features by deep learning, we use an autoencoder with one encoding layer and one decoding layer. The activation function for both layers is the hyperbolic tangent (“tanh”), and the reconstruction loss is defined as the mean squared error (MSE) between the input **x** and the reconstructed output 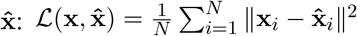:, where **z** = tanh(**xw**_enc_ + **b**_enc_) is the encoded latent representation, and 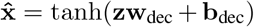 is the reconstructed data. Here, (**w**_enc_, **b**_enc_, **w**_dec_, **b**_dec_) are the parameters of the encoder and decoder layers, respectively. By learning these parameters, we could minimize *L*, and thus get the representation for latent features **z**.

## 3 Simulation Study

### 3.1 Simulation Settings

We conducted extensive simulation studies to evaluate the performance of our method in terms of detecting truly associated risk factors, identifying effect modifications, and predicting the outcome. We considered various sample sizes *N* = 200, 400, 1000, 2000, 4000, 6000, with the proportion of cases set to 50%. We set the risk factor dimension at *p* = 100, the underlying factor dimension at *K* = 10, and the number of factors extracted in the method to *r* = 2, 5, 8, 10. These relationships between covariates and factors are modeled differently for data generation as specified below. This relationship could be either linear or nonlinear. The results show that the existing method, which cannot accommodate the factor effect, yields greater biases for both outcome prediction and parameter estimation. By varying choices of N and K, we thoroughly investigated the finite sample performance of our proposed method under a variety of simulation settings, characterized by different factor-covariate associations. The random K-dimensional factors **z** follow the standard normal distribution, and the random error *u*_*i*_ follows a normal distribution with mean 0 and variance 0.2. We considered two scenarios considering both the linear and nonlinear relationship between **x** and **z**. For the linear relationship, we generated **x** from the mechanism: **x**_*i*_ = Λ**z**_*i*_ + **u**_*i*_. For Λ, we considered the following two cases: (a) Λ = [*l*_1_ … *l*_1_], with 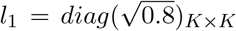. (b) Λ is generated by optimizing a random matrix to approximate a target matrix *A*, which is constructed using a random orthogonal matrix *Q* and a diagonal matrix of exponentially distributed eigenvalues. The optimization minimizes the Frobenius norm of the difference between ΛΛ^*T*^ and *A*, and the final Λ is standardized so that the sum of squares of each column is 0.8. The objective function is: min_Λ_ ‖ ΛΛ^*T*^ − *A* ‖_*F*_, where *A* = *Q*Λ_diag_*Q*^*T*^.

For nonlinear relationships, we also considered two cases: (c) Using **x**, which was previously generated by case (b), we fitted a one-layer autoencoder model and extracted the factors to represent the latent features **z**. (d) The matrix **x** is generated using a generalized additive model, where for each column a unique sequence of sine or cosine functions is applied to random variables **z**, with sine and cosine coefficients drawn from uniform distribution: *U* (0.5, 1.5). The randomness in the function selection and the factors add variability to the model. The generation algorithm is shown below:

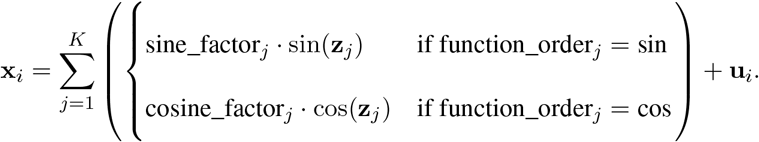

In (a), we aimed to simulate an equal factor relationship, which could be a useful representation for solving batch effects. In (b), we considered a more generalized setting where factors have significantly different effects. In (c), we simulated an autoencoder mapping **x** to **z** then back to **x**. In (d), a generalized additive models was used to simulate nonlinear relationships. We generated the outcomes using this equation: 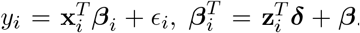. Due to sparsity requirements, we allowed only 10% entries of ***β***_;*i*_to be nonzero. Additionally, **δ, *β*** and ***ϵ*** are designed so that 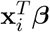 and 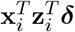 each explain 25% of the variability in y, with ***ϵ*** explaining the other 50%.

In the following, we compare the performance of LF-VCR, LF-VCR (ae) with those commonly used approaches: (i) Lasso; (ii) Elastic Net, which ignores any possible factor effects. All results are summarized from the corresponding test set. In the following, we present the result with *r* = 5. Other results are presented in the sensitivity section in the Supplement.

### 3.2 Simulation Results

We evaluated the performance of our proposed methods, LF-VCR and LF-VCR (ae), in comparison to Lasso and Elastic Net across various settings. Below, we present our results in two sections: the first is for setting (a) and (b), where contributing latent features have linear effects on ***β***_*i*_; the second is for setting (c) and (d), where contributing latent features have nonlinear effects on ***β***_*i*_. We summarize the key findings in predictive accuracy, parameter estimation, and factor robustness under different settings. In Table 1, results on the prediction of outcomes are summarized by using *R*^2^ comparison between LF-VCR, LF-VCR (ae) and Lasso, Elastic Net to help reveal the role of factors in outcome prediction. In Table 2, estimation results for the MSE for 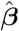 are compared across the four comparison methods to reveal parameter estimation accuracy. The visualizations for both tables are presented in Figure 1.

**Table 1:** Simulation results under different settings with varying sample sizes (*N* = 200, 400, 1000, 2000, 4000, 6000) and a fixed risk factor dimension (*p* = 100). Prediction R-squared (*R*^2^) measured from the test set from LF-VCR, LF-VCR (ae), Lasso, Elastic Net was reported under four different settings: *(a) contributing latent features have identical linear effects on* ***β***_*i*_, *(b) contributing latent features have descending linear effects on* ***β***_*i*_, *(c) autoencoder model with nonlinear latent feature contributions, and (d) latent features contribute in a generalized additive model (GAM) with nonlinear effects on* ***β***_*i*_.

| N | Setting | Prediction $R^2$ | | | |
| --- | --- | --- | --- | --- | --- |
|  |  | LF-VCR | LF-VCR (ae) | Lasso | Elastic Net |
| 200 | (a) | 0.1479 | 0.1383 | 0.1159 | 0.1252 |
|  | (b) | 0.2167 | 0.1990 | 0.1361 | 0.1366 |
|  | (c) | 0.3080 | 0.3561 | 0.1826 | 0.1854 |
|  | (d) | 0.2993 | 0.3188 | 0.2803 | 0.2834 |
| 400 | (a) | 0.2281 | 0.2142 | 0.1445 | 0.1499 |
|  | (b) | 0.2718 | 0.2651 | 0.1579 | 0.1592 |
|  | (c) | 0.3902 | 0.4206 | 0.2052 | 0.2081 |
|  | (d) | 0.3403 | 0.3471 | 0.3108 | 0.3134 |
| 1000 | (a) | 0.3406 | 0.3341 | 0.2037 | 0.2037 |
|  | (b) | 0.3217 | 0.3181 | 0.1711 | 0.1718 |
|  | (c) | 0.4416 | 0.4565 | 0.2145 | 0.2163 |
|  | (d) | 0.3774 | 0.3747 | 0.3336 | 0.3351 |
| 2000 | (a) | 0.3808 | 0.3823 | 0.2194 | 0.2191 |
|  | (b) | 0.3524 | 0.3506 | 0.1915 | 0.1919 |
|  | (c) | 0.4708 | 0.4809 | 0.2238 | 0.2248 |
|  | (d) | 0.3934 | 0.3908 | 0.3446 | 0.3453 |
| 4000 | (a) | 0.4142 | 0.4110 | 0.2363 | 0.2357 |
|  | (b) | 0.3670 | 0.3665 | 0.1990 | 0.1980 |
|  | (c) | 0.4877 | 0.4980 | 0.2353 | 0.2354 |
|  | (d) | 0.4095 | 0.4093 | 0.3537 | 0.3538 |
| 6000 | (a) | 0.4246 | 0.4267 | 0.2397 | 0.2391 |
|  | (b) | 0.3730 | 0.3719 | 0.2001 | 0.2001 |
|  | (c) | 0.4940 | 0.5036 | 0.2393 | 0.2392 |
|  | (d) | 0.4172 | 0.4177 | 0.3597 | 0.3600 |

**Table 2:** Simulation results under different settings with varying sample sizes (*N* = 200, 400, 1000, 2000, 4000, 6000) and a fixed risk factor dimension (*p* = 100). Mean Squared Error (MSE) of 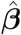 measured from the test set from LF-VCR, LF-VCR (ae), Lasso, Elastic four different settings: *(a) contributing latent features have identical linear effects on* ***β***_*i*_, *(b) contributing latent features have descending linear effects on* ***β***_*i*_, *(c) autoencoder model with nonlinear latent feature contributions, and (d) latent features contribute in a GAM with nonlinear effects on* ***β***_*i*_.

| N | Setting | MSE for $\hat{\beta}$ | | | |
| --- | --- | --- | --- | --- | --- |
|  |  | LF-VCR | LF-VCR (ae) | Lasso | Elastic Net |
| 200 | (a) | 0.4990 | 0.5050 | 0.5165 | 0.5093 |
|  | (b) | 0.3178 | 0.3572 | 0.3124 | 0.3070 |
|  | (c) | 0.3022 | 0.3142 | 0.3065 | 0.3057 |
|  | (d) | 0.1028 | 0.0843 | 0.1486 | 0.1488 |
| 400 | (a) | 0.4251 | 0.4265 | 0.4264 | 0.4234 |
|  | (b) | 0.2801 | 0.2792 | 0.2584 | 0.2466 |
|  | (c) | 0.2438 | 0.2431 | 0.2624 | 0.2456 |
|  | (d) | 0.0815 | 0.0678 | 0.1155 | 0.1049 |
| 1000 | (a) | 0.2963 | 0.2982 | 0.3110 | 0.3110 |
|  | (b) | 0.2145 | 0.2209 | 0.2167 | 0.2137 |
|  | (c) | 0.1866 | 0.1742 | 0.2092 | 0.2005 |
|  | (d) | 0.0690 | 0.0630 | 0.1002 | 0.0940 |
| 2000 | (a) | 0.2155 | 0.2191 | 0.2456 | 0.2482 |
|  | (b) | 0.1781 | 0.1826 | 0.1915 | 0.1907 |
|  | (c) | 0.1423 | 0.1292 | 0.1717 | 0.1682 |
|  | (d) | 0.0617 | 0.0624 | 0.0963 | 0.0964 |
| 4000 | (a) | 0.1580 | 0.1610 | 0.2050 | 0.2080 |
|  | (b) | 0.1471 | 0.1520 | 0.1660 | 0.1670 |
|  | (c) | 0.1120 | 0.0880 | 0.1427 | 0.1431 |
|  | (d) | 0.0580 | 0.0630 | 0.0990 | 0.0960 |
| 6000 | (a) | 0.1350 | 0.1370 | 0.1910 | 0.1940 |
|  | (b) | 0.1270 | 0.1340 | 0.1530 | 0.1530 |
|  | (c) | 0.0960 | 0.0694 | 0.1290 | 0.1310 |
|  | (d) | 0.0571 | 0.0632 | 0.0990 | 0.0980 |

**Figure 1:**
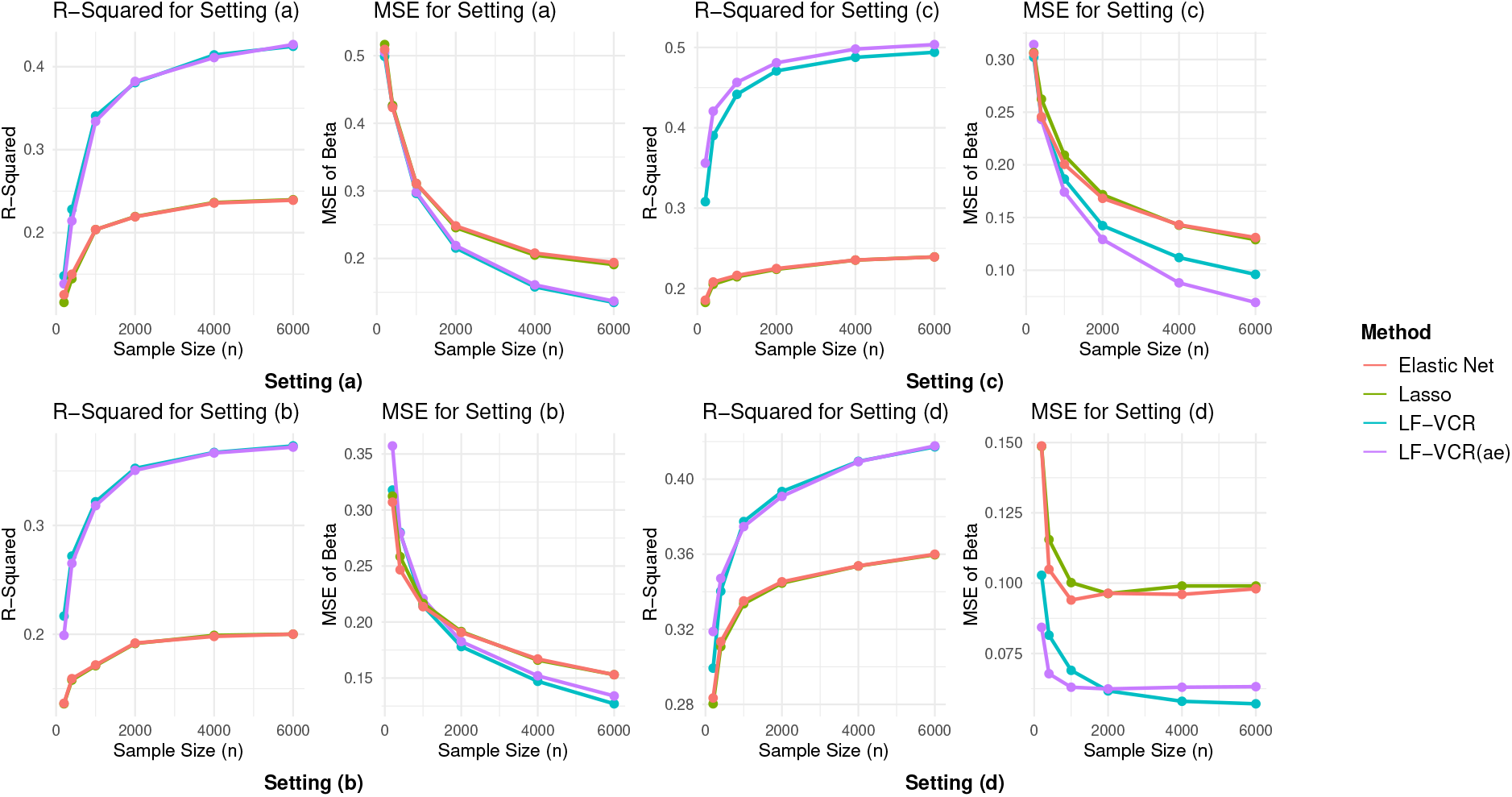
Simulation results illustrating the prediction *R*^2^ and Mean Squared Error (MSE) for 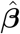 across four methods: LF-VCR, LF-VCR (ae), Lasso, and Elastic Net. The analysis was conducted under four different simulation settings with varying sample sizes (*N* = 200, 400, 1000, 2000, 4000, 6000) and a fixed risk factor dimension (*p* = 100). *The sparsity of risk factors with nonzero effects is set to 10%. We consider four different settings in terms of how* ***β***_*i*_ *varies by the latent factors: (a) contributing latent features have equal linear effects on* ***β***_*i*_, *(b) contributing latent features have descending linear effects on* ***β***_*i*_, *(c) autoencoder model with nonlinear latent feature contributions, and (d) latent features contribute in a generalized additive model (GAM) with nonlinear effects on* ***β***_*i*_.

**Figure 2:**
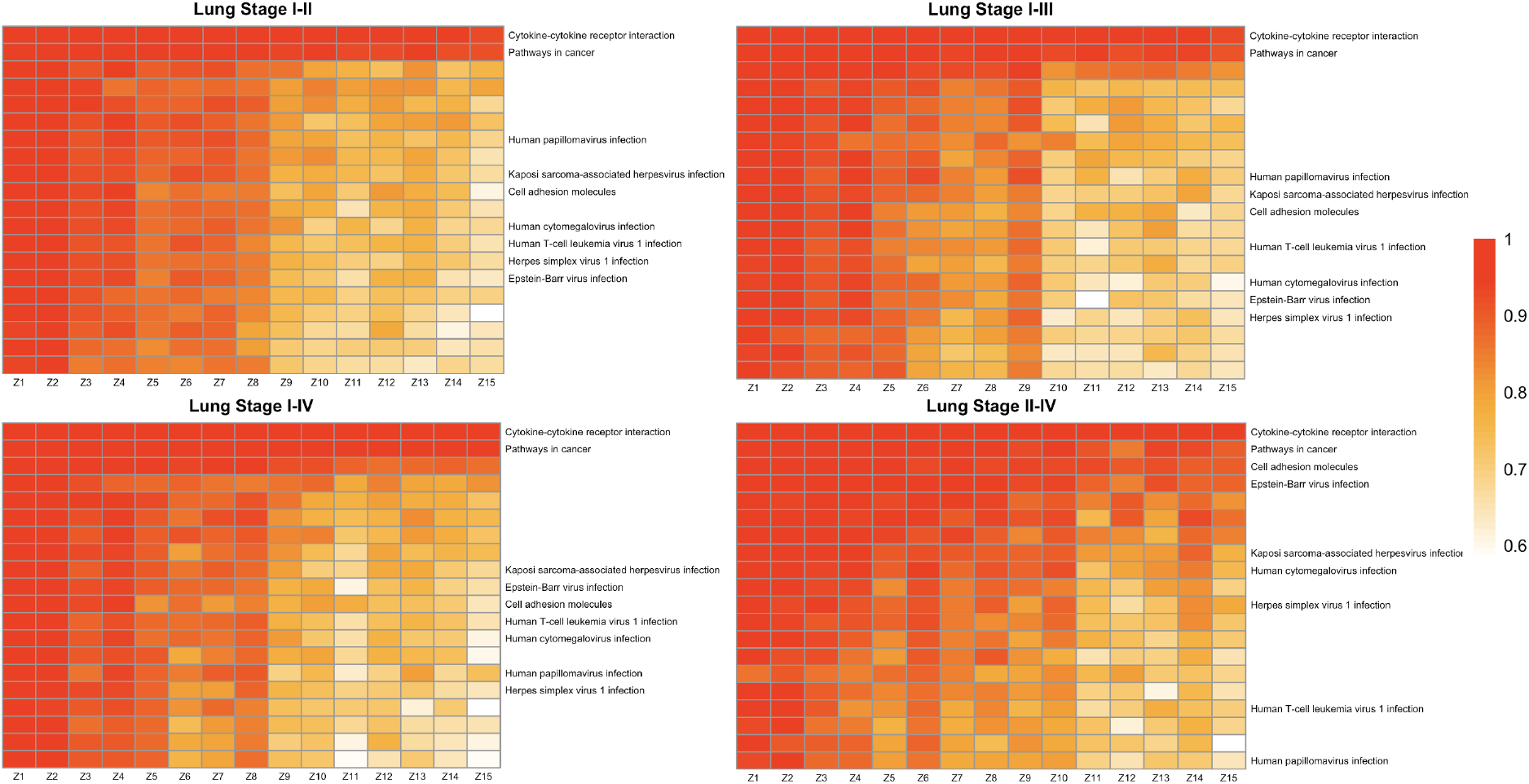
Factor-Gene Pathway correlations in different lung cancer stage pairs. The graph presents the top 20 pathways for each pair of samples, while labeling only the 9 pathways that are common across the datasets. *The x-axis (Z1 to Z15) represents the 15 extracted factors and the y-axis represents different gene pathways. For each gene pathway, linear regression was used to fit the genes in the pathway against all factors, and the corresponding R-squared were presented*

#### 3.2.1 Simulation Studies Assuming Linear Latent Features

We first examined the method performance under settings (a) and (b), where latent features present a linear effect modification on risk outcomes. For setting (a), the effect modification is the same across all contributing latent features, and for setting (b): the effect modification has a descending trend across different latent features. Next, we evaluated predictive performance by analyzing the MSE, and *R*^2^ for outcome prediction. LF-VCR and LF-VCR (ae) consistently achieved lower MSEs compared to Lasso and Elastic Net, particularly as the sample size *N* increased. For smaller samples (*N* = 200, 400), the differences in MSE were minimal across all methods. However, as *N* increased, the superiority of LF-VCR and LF-VCR (ae) became more pronounced. For *N* = 4000 and *N* = 6000, the MSEs for LF-VCR and LF-VCR (ae) stabilized at approximately 0.6, whereas Lasso and Elastic Net remained at approximately 0.8. Under the equal factor size scenario (a), LF-VCR and LF-VCR (ae) initially exhibited higher MSEs for smaller sample sizes compared to the descending factor size scenario (b). However, their MSEs decreased more rapidly as the sample size increased, ultimately achieving lower values. As anticipated, Lasso and Elastic Net produced similar results, and a comparable trend was observed for LF-VCR and LF-VCR (ae). This resemblance likely stems from the assumed linear relationships during feature generation, which may have led LF-VCR and LF-VCR (ae) to extract similar features. In terms of *R*^2^, LF-VCR and LF-VCR (ae) consistently outperformed the other methods. Although the differences in *R*^2^ were modest for smaller sample sizes, the gap widened as *N* increased. For larger *N, R*^2^ for LF-VCR and LF-VCR (ae) approached 0.375, while Lasso and Elastic Net maintained values around 0.23. These findings indicate that LF-VCR and LF-VCR (ae) exhibit superior predictive accuracy compared to Lasso and Elastic Net across various settings.

We further evaluated the performance of the models in terms of parameter estimations by examining the MSE for 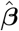, as well as the true positive rate (TPR) and false positive rate (FPR) of identifying truly associated features. For smaller sample sizes (*N* = 200 or 400), the MSE for 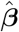 was comparable across all methods. However, as *N* increased, LF-VCR and LF-VCR (ae) demonstrated a more pronounced decline in MSE for 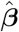 compared to Lasso and Elastic Net, ultimately achieving significantly lower values for *N* ≥ 2000. Detailed results are summarized in Tables S1 to S6 (Supplementary Materials). We analyzed how the predictive power varied with different values of *r*, the number of latent factors extracted. Both LF-VCR and LF-VCR (ae) exhibit a consistent trend with respect to *r*. Generally, the MSE for *y* and the MSE for 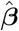 decrease as *r* increases. However, further increases in *r* beyond 5 result in negligible reductions in MSE.

#### 3.2.2 Simulation Studies Assuming Nonlinear Latent Features

In this section, we examine our results for settings (c) and (d), where latent features present a nonlinear effect modification on risk outcomes. For setting (c), the effect modification is generated through an autoencoder process, and for setting (d) the effect modification is generated through a generalized additive model with sin, cos function. The results under nonlinear settings were similar to those in the linear case. LF-VCR (ae) outperformed LF-VCR at smaller sample sizes, achieving lower MSEs and higher *R*^2^. As *N* increased, the performance gap between LF-VCR and LF-VCR (ae) narrowed. For larger *N*, both methods stabilized at similar levels of performance. Compared to LF-VCR (ae), LF-VCR was more robust under most settings when the sample size becomes larger. Regarding parameter estimation, MSE for 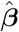 showed a similar trend. For smaller *N*, all four methods demonstrated similar accuracy. LF-VCR (ae) exhibited the fastest decline, dropping below 0.1 for *N* = 6000, while LF-VCR showed a slower but significant improvement compared to Lasso and Elastic Net.

In both linear and nonlinear scenarios, LF-VCR and LF-VCR (ae) were computationally more intensive than Lasso and Elastic Net, owing to factor extraction and interaction modeling. Nevertheless, the additional computational burden was justified by the significant improvements in predictive accuracy and parameter estimation.

## 4 A real data example

In lung cancer diagnosis, extensive research has been conducted utilizing gene expression profiling to identify pathways associated with lung cancer development and progression, for example, (Dancik and Theodorescu (2014);Chang et al. (2015);Cai and Jiang (2014)). In this application, we aim to predict different lung cancer stages and identify important risk factors and corresponding gene pathways. These findings could provide insights into the underlying genetic mechanisms of lung cancer progression.

We applied LF-VCR, LF-VCR (ae), as well as Lasso and Elastic Net regressions, the two existing methods also considered in the simulation studies, to a gene expression dataset for lung cancer patients in The Cancer Genome Atlas (TCGA) Program (Network (2014)). The dataset used in this analysis was sourced from the Lung Cancer Explorer (LCE) (Quantitative Biomedical Research Center, 2019) and underwent standard quality control procedures in a previous study Jin and Wang (2022). Zero expression values were replaced with the overall minimum value, and all data were subsequently log_2_-transformed to approximate a normal distribution across all lung cancer stages (Cai and Jiang, 2014; Cai et al., 2019). Gene expression levels for 20,429 genes were measured in lung cancer tissue samples obtained from 513 patients, distributed as follows: *N*_1_ = 278, *N*_2_ = 124, *N*_3_ = 84, and *N*_4_ = 27 for stages I, II, III, and IV, respectively. Additionally, *N*_0_ = 59 normal tissue samples were collected from these patients. To avoid dependency when comparing tumor tissues with normal tissues, tumor samples from these 59 patients were excluded from the analysis.

We considered *H* = 206 common human pathways related to signaling, metabolic, and human diseases from the KEGG database (Kanehisa et al., 2019; Kanehisa and Goto, 2000). Based on differentially expressed genes in these pathways, we constructed four comparison groups: Stage I versus Stage II, Stage I versus Stage III, Stage I versus Stage IV, and Stage II versus Stage IV. For the comparison between stage I and stage II lung cancer, we have sample sizes for the two groups *N*_1_ = 278; *N*_2_ = 124, with a total of *p* = 1, 862 genes within differentially expressed pathways previously detectedJin and Wang (2022). For stage I versus stage III, we have sample sizes for the two groups *N*_1_ = 278; *N*_3_ = 84, with *p* = 2, 857 genes. For stage I versus stage IV, we have sample sizes for the two groups *N*_1_ = 278; *N*_4_ = 27, with *p* = 2, 036 genes. For stage II versus stage IV, we have sample sizes for the two groups *N*_2_ = 124; *N*_4_ = 27, with *p* = 749 genes within differentially expressed pathways.

We randomly divided the data into training (80%, for model development) and testing (20%, for model evaluation) sets, with a 5-fold cross-validation. The reported results were obtained as the average across 100 repeats of the 5-fold cross-validation. Detailed results are summarized in Table 3. Our method had a 6% increase in (AUC-0.5) for Stage I versus Stage II and a 10% increase for Stage II versus Stage IV compared to lasso and elastic net. Notably, the improvement was even more pronounced in the other two comparisons, with a 61% increase in (AUC-0.5) for Stage I versus Stage III and a remarkable 118% increase for Stage I versus Stage IV. This significant improvement was achieved not by incorporating additional information but by fully leveraging the existing data. The more pronounced results for Stage I versus Stage III and Stage I versus Stage IV, compared to Stage I versus Stage II, may be attributed to the presence of more differentially expressed genes across distinct gene pathways. This facilitated the extracted factors in capturing more precise gene pathway information. By incorporating this information through a group structure, our method was able to achieve superior classification results.

**Table 3:** Comparison of AUC between different pairs of lung cancer stages. *Bold font indicates the highest AUC for the classification of each pair of lung cancer stages*.

| Lung Cancer Stage | AUC |  |  |  |
| --- | --- | --- | --- | --- |
|  | LF-VCR | LF-VCR (ae) | Lasso | Elastic Net |
| Stage I vs. II | **0.5889** | 0.5888 | 0.5836 | 0.5854 |
| Stage I vs. III | **0.6305** | 0.6276 | 0.5810 | 0.5992 |
| Stage I vs. IV | 0.5848 | **0.5851** | 0.5388 | 0.5418 |
| Stage II vs. IV | **0.5547** | 0.5531 | 0.5495 | 0.5435 |

To better interpret the hidden factors, we analyzed the relationship between gene pathways and factors by fitting a linear regression for each factor against differentially expressed genes in each pathways. Across all four pairs, out of the first 20 pathways showing high correlation to all the factors we identified 9 common gene pathways. Many of these pathways highlight mechanisms like inflammation, immune modulation, and cellular signaling, which are relevant to lung cancer biology. Alterations in these processes can contribute to tumor initiation, growth, immune evasion, and metastasis. The extracted factors highly represented the significant pathways for lung cancer expression, leading to significant improvement in prediction accuracy without integrating any other information.

## 5 Discussion

We proposed LF-VCR, a novel varying coefficient regression framework to model associations between risk factors (exposures) and health outcomes by accounting for potential effect modification from latent features of the risk factors. Our method allows the effects of some risk factors to vary as a function of latent features extracted through either statistical factor analysis (e.g., POET) or deep learning (e.g., autoencoders), which are hypothesized to capture unobserved environmental factors that affect the exposure-outcome associations. The key feature of LF-VCR is the varying-by-latent feature model coefficients, which, in the absence of data for other population characteristics that affect the exposure-outcome associations, can capture residual effect modifications by accounting for interactions between risk factors and their latent features, which are often overlooked in conventional approaches. It can help better identify true risk factors and critical interaction effects, improving our understanding of the exposure-outcome relationships. We applied sparse-group-lasso penalties to induce sparsity within and across interaction groups, ensuring robustness and interoperability of the detection of effect modification by risk factors, improving variable selection for identifying risk factors truly associated with the outcome, and empowering prediction of the outcome. This has been illustrated by extensive simulations, where our method outperforms standard penalized regressions, including lasso and elastic net, without considering the varying-by-latent-feature, in terms of model coefficient estimation and predictive accuracy on the outcome in various high-dimensional data scenarios. Our simulation results have also shown that the deep learning autoencoder can outperform the linear factor model in latent feature extraction in the presence of complex, nonlinear structures, and this advantage still appears with a relatively small training sample size. Its application to a TCGA dataset demonstrates its superiority in classifying lung cancer stages based on gene expression data compared to alternative approaches with fixed model coefficients and highlights the method’s capability to address real-world challenges in the high-dimensional setting.

Our study has several limitations. First, our current model requires substantial computational time due to the dual steps of feature extraction and model fitting, which can be resource-intensive. Solutions to this computational issue are needed, especially for applications in large N (e.g., *N >* 100, 000, large p (e.g., *p >* 100, 000) settings, which are common in large-scale genetic and genomic studies. To reduce computational burden, we can optimize the feature extraction step by implementing it with a selected subset of high-dimensional risk factors, such as generating genetic ancestry descriptors with only ancestrally informative genetic variants in genetic risk prediction models Qu et al. (2019); Alladio et al. (2022). Further dimension reduction techniques can be considered to reduce the computational burden for the model fitting step as well. Second, the model only considers linear effect modifications by latent features, whichn basically adds linear interaction terms between risk factors and their latent features, whereas these modifications may follow a functional form or could be better captured using machine learning techniques. Future work is needed to explore these alternative approaches. Lastly, interpreting the extracted latent features can be challenging, as their underlying meaning may not always be clear or directly relatable to the observed risk factors.

Our current model only accounts for linear effect modification, but it can be extended to accommodate nonlinear interactions by incorporating ensemble models or neural networks. Such enhancements could further improve predictive performance for those tasks focusing on prediction. For inference-driven objectives, our method could incorporate debiased group lasso, which would allow more robust and valid inference and better identification of disease-associated biomarkers, contributing to the understanding of disease mechanisms and new drug target identification. Additionally, a higher-dimensional version of LF-VCR could be adapted to address multi-ancestry analysis in GWAS, broadening its application to genetic studies. Looking forward, these advancements have the potential to significantly aid precision medicine by enabling personalized predictions and insights. Despite these opportunities for extension, we continue to value the simplicity of our current model setup, as it offers the advantage of ease of implementation and interoperability.

## Data Availability

The data that support the findings in this paper are available to download from the Lung Cancer Explorer (LCE) at https://lce.biohpc.swmed.edu/lungcancer/.

