## Supplementary Materials for "Identifying Effect Modification of Latent Population Characteristics on Risk Factors with a Sparse Varying Coefficient Regression"

### Supplementary Materials for "Identifying Effect Modification of Latent Population Characteristics on Risk Factors with a Sparse Latent Factor Regression"

Ruofan Wang[1], Lei Fang[2], Yue Wang[3], and Jin Jin[1]

[1]Department of Biostatistics, Epidemiology and Informatics, University of Pennsylvania, Philadelphia, PA 19104, USA
[2]Department of Biostatistics and Data Science, University of Minnesota Twin Cities, Minneapolis, MN 55455, USA
[3]Department of Biostatistics and Informatics, Colorado School of Public Health, 13001 E. 17th Place, Aurora, CO 80045, USA

We report additional simulation results from Section 3 in the Supplementary Materials.

Table S1: *Results of simulations in Section 3: summarized from 100 Simulations under setting (a) and (b) with* **n=200, p=100.**

| **Setting** | | | **r=2** | **r=5** | **r=8** | **r=10** |
|---|---|---|---|---|---|---|
| Setting(a) $\mathbf{x}_i = \Lambda \mathbf{z}_i + \mathbf{u}_i$ $\Lambda = [\mathbf{l_1}, \dots, \mathbf{l_1}]$ $\mathbf{l}_1 = diag(\sqrt{0.8})_{K\times K}$ | VF | MSE | 0.9363 | 0.9079 | 0.9142 | 0.9128 |
| | | R-square | 0.1297 | 0.1479 | 0.1572 | 0.1488 |
| | | MSE for beta | 0.5063 | 0.499 | 0.5208 | 0.4991 |
| | | Main Effect TPR | 0.287 | 0.311 | 0.349 | 0.33 |
| | | Main Effect FPR | 0.112 | 0.1245 | 0.1367 | 0.1264 |
| | | Interaction TPR | 0.0793 | 0.0863 | 0.1093 | 0.0984 |
| | | Interaction FPR | 0.0754 | 0.3345 | 0.7385 | 0.8541 |
| | VF-ae | MSE | 0.9568 | 0.9256 | 0.9059 | 0.9358 |
| | | R-square | 0.1098 | 0.1383 | 0.1468 | 0.1402 |
| | | MSE for beta | 0.5202 | 0.5049 | 0.4906 | 0.5184 |
| | | Main Effect TPR | 0.264 | 0.297 | 0.307 | 0.304 |
| | | Main Effect FPR | 0.108 | 0.1195 | 0.118 | 0.1282 |
| | | Interaction TPR | 0.0684 | 0.0901 | 0.0961 | 0.1011 |
| | | Interaction FPR | 0.0658 | 0.3469 | 0.6492 | 0.8779 |
| Setting(b) $\mathbf{x}_i = \Lambda \mathbf{z}_i + \mathbf{u}_i$ $\Lambda = \min_\Lambda \left\|\Lambda\Lambda^T - A\right\|_F$ $A = Q\Lambda_{\text{diag}}Q^T$ | VF | MSE | 0.8743 | 0.8024 | 0.8126 | 0.8252 |
| | | R-square | 0.154 | 0.2167 | 0.1992 | 0.1875 |
| | | MSE for beta | 0.3128 | 0.3178 | 0.2961 | 0.2865 |
| | | Main Effect TPR | 0.195 | 0.235 | 0.225 | 0.218 |
| | | Main Effect FPR | 0.06 | 0.0688 | 0.069 | 0.0626 |
| | | Interaction TPR | 0.0587 | 0.273 | 0.4372 | 0.4793 |
| | VF-ae | MSE | 0.8807 | 0.8249 | 0.83 | 0.8326 |
| | | R-square | 0.1491 | 0.199 | 0.2004 | 0.1985 |
| | | MSE for beta | 0.3266 | 0.3572 | 0.3567 | 0.3607 |
| | | Main Effect TPR | 0.202 | 0.243 | 0.239 | 0.253 |
| | | Main Effect FPR | 0.065 | 0.083 | 0.081 | 0.0848 |
| | | Interaction TPR | 0.0721 | 0.2931 | 0.4381 | 0.5673 |

Table S2: *Results of simulations in Section 3: summarized from 100 Simulations under setting (a) and (b) with* **n=400, p=100.**

| **Setting** | | | **r=2** | **r=5** | **r=8** | **r=10** |
|---|---|---|---|---|---|---|
| Setting(a) $\mathbf{x}_i = \Lambda \mathbf{z}_i + \mathbf{u}_i$ $\Lambda = [\mathbf{l_1}, \ldots, \mathbf{l_1}]$ $\mathbf{l}_1 = \text{diag}(\sqrt{0.8})_{K\times K}$ | LF-VCR | MSE | 0.8293 | 0.7784 | 0.7619 | 0.7541 |
| | | R-square | 0.1826 | 0.2281 | 0.2466 | 0.2533 |
| | | MSE for beta | 0.4472 | 0.4251 | 0.4230 | 0.4188 |
| | | Main Effect TPR | 0.556 | 0.595 | 0.615 | 0.619 |
| | | Main Effect FPR | 0.1641 | 0.1725 | 0.1805 | 0.1837 |
| | | Interaction TPR | 0.1346 | 0.1686 | 0.1913 | 0.1915 |
| | | Interaction FPR | 0.1294 | 0.6506 | 1.2937 | 1.6586 |
| | LF-VCR (ae) | MSE | 0.8421 | 0.7920 | 0.7579 | 0.7582 |
| | | R-square | 0.1654 | 0.2142 | 0.2453 | 0.2475 |
| | | MSE for beta | 0.4336 | 0.4265 | 0.4205 | 0.4211 |
| | | Main Effect TPR | 0.433 | 0.561 | 0.598 | 0.607 |
| | | Main Effect FPR | 0.1553 | 0.1651 | 0.1768 | 0.1824 |
| | | Interaction TPR | 0.1239 | 0.1606 | 0.1834 | 0.1866 |
| | | Interaction FPR | 0.1211 | 0.6214 | 1.2369 | 1.6205 |
| Setting(b) $\mathbf{x}_i = \Lambda \mathbf{z}_i + \mathbf{u}_i$ $\Lambda = \min_\Lambda \left\|\Lambda\Lambda^T - A\right\|_F$ $A = Q\Lambda_{\text{diag}}Q^T$ | LF-VCR | MSE | 0.8409 | 0.7496 | 0.7603 | 0.7694 |
| | | R-square | 0.1824 | 0.2718 | 0.2581 | 0.2501 |
| | | MSE for beta | 0.2725 | 0.2801 | 0.2543 | 0.2511 |
| | | Main Effect TPR | 0.338 | 0.379 | 0.363 | 0.354 |
| | | Main Effect FPR | 0.0783 | 0.0966 | 0.0803 | 0.0752 |
| | | Interaction TPR | 0.1041 | 0.4310 | 0.6675 | 0.7847 |
| | LF-VCR (ae) | MSE | 0.8440 | 0.7536 | 0.7491 | 0.7539 |
| | | R-square | 0.1805 | 0.2651 | 0.2689 | 0.2647 |
| | | MSE for beta | 0.2637 | 0.2792 | 0.2703 | 0.2727 |
| | | Main Effect TPR | 0.320 | 0.377 | 0.377 | 0.383 |
| | | Main Effect FPR | 0.0682 | 0.1030 | 0.0943 | 0.1000 |
| | | Interaction TPR | 0.0903 | 0.4043 | 0.5719 | 0.7585 |

Table S3: *Results of simulations in Section 3: summarized from 100 Simulations under setting (a) and (b) with n=1000, p=100.*

| **n, p** | | | | **r=2** | **r=5** | **r=8** | **r=10** |
|---|---|---|---|---|---|---|---|
| 1000, 100 | (a) | LF-VCR | MSE | 0.7325 | 0.6587 | 0.6245 | 0.607 |
| | | | R-square | 0.2699 | 0.3406 | 0.3747 | 0.3922 |
| | | | MSE for beta | 0.3225 | 0.2963 | 0.2749 | 0.2559 |
| | | | Main Effect TPR | 0.897 | 0.925 | 0.93 | 0.933 |
| | | | Main Effect FPR | 0.2118 | 0.2144 | 0.2235 | 0.227 |
| | | | Interaction TPR | 0.2393 | 0.3162 | 0.329 | 0.3296 |
| | | | Interaction FPR | 0.2304 | 1.2201 | 2.2255 | 2.8657 |
| | | LF-VCR (ae) | MSE | 0.7419 | 0.6651 | 0.6232 | 0.6091 |
| | | | R-square | 0.2599 | 0.3341 | 0.3758 | 0.3901 |
| | | | MSE for beta | 0.3203 | 0.2983 | 0.2763 | 0.2581 |
| | | | Main Effect TPR | 0.889 | 0.904 | 0.928 | 0.93 |
| | | | Main Effect FPR | 0.205 | 0.2162 | 0.221 | 0.2251 |
| | | | Interaction TPR | 0.2292 | 0.3285 | 0.3492 | 0.3395 |
| | | | Interaction FPR | 0.2217 | 1.2672 | 2.3466 | 2.9492 |
| | (b) | LF-VCR | MSE | 0.7761 | 0.6688 | 0.676 | 0.6793 |
| | | | R-square | 0.2132 | 0.3216 | 0.3151 | 0.3115 |
| | | | MSE for beta | 0.2346 | 0.2145 | 0.2143 | 0.2116 |
| | | | Main Effect TPR | 0.524 | 0.592 | 0.59 | 0.589 |
| | | | Main Effect FPR | 0.102 | 0.1108 | 0.108 | 0.1073 |
| | | | Interaction TPR | 0.1485 | 0.5581 | 0.9359 | 1.1707 |
| | | LF-VCR (ae) | MSE | 0.7759 | 0.6722 | 0.6696 | 0.6723 |
| | | | R-square | 0.2128 | 0.318 | 0.3205 | 0.3175 |
| | | | MSE for beta | 0.2328 | 0.2209 | 0.219 | 0.2244 |
| | | | Main Effect TPR | 0.524 | 0.605 | 0.606 | 0.609 |
| | | | Main Effect FPR | 0.095 | 0.122 | 0.1224 | 0.1296 |
| | | | Interaction TPR | 0.147 | 0.5258 | 0.8021 | 1.0624 |

Table S4: *Results of simulations in Section 3: summarized from 100 simulations under setting (a) and (b) with* **n=1000, p=100.**

| **Setting** | | | **r=2** | **r=5** | **r=8** | **r=10** |
|---|---|---|---|---|---|---|
| Setting (a)<br>$\mathbf{x}_i = \Lambda \mathbf{z}_i + \mathbf{u}_i$<br>$\Lambda = [\mathbf{l_1}, \ldots, \mathbf{l_1}]$<br>$\mathbf{l}_1 = \text{diag}(\sqrt{0.8})_{K \times K}$ | LF-VCR | MSE | 0.7325 | 0.6587 | 0.6245 | 0.6070 |
| | | R-square | 0.2699 | 0.3406 | 0.3747 | 0.3922 |
| | | MSE for beta | 0.3225 | 0.2963 | 0.2749 | 0.2559 |
| | | Main Effect TPR | 0.897 | 0.925 | 0.930 | 0.933 |
| | | Main Effect FPR | 0.2118 | 0.2144 | 0.2235 | 0.2270 |
| | | Interaction TPR | 0.2393 | 0.3162 | 0.3290 | 0.3296 |
| | | Interaction FPR | 0.2304 | 1.2201 | 2.2255 | 2.8657 |
| | LF-VCR (ae) | MSE | 0.7419 | 0.6651 | 0.6232 | 0.6091 |
| | | R-square | 0.2599 | 0.3341 | 0.3758 | 0.3901 |
| | | MSE for beta | 0.3203 | 0.2983 | 0.2763 | 0.2581 |
| | | Main Effect TPR | 0.889 | 0.904 | 0.928 | 0.930 |
| | | Main Effect FPR | 0.2050 | 0.2162 | 0.2210 | 0.2251 |
| | | Interaction TPR | 0.2292 | 0.3285 | 0.3492 | 0.3395 |
| | | Interaction FPR | 0.2217 | 1.2672 | 2.3466 | 2.9492 |
| Setting (b)<br>$\mathbf{x}_i = \Lambda \mathbf{z}_i + \mathbf{u}_i$<br>$\Lambda = \min_\Lambda \left\|\Lambda\Lambda^T - A\right\|_F$<br>$A = Q\Lambda_{\text{diag}}Q^T$ | LF-VCR | MSE | 0.7761 | 0.6688 | 0.6760 | 0.6793 |
| | | R-square | 0.2132 | 0.3216 | 0.3151 | 0.3115 |
| | | MSE for beta | 0.2346 | 0.2145 | 0.2143 | 0.2116 |
| | | Main Effect TPR | 0.524 | 0.592 | 0.590 | 0.589 |
| | | Main Effect FPR | 0.1020 | 0.1108 | 0.1080 | 0.1073 |
| | | Interaction TPR | 0.1485 | 0.5581 | 0.9359 | 1.1707 |
| | LF-VCR (ae) | MSE | 0.7759 | 0.6722 | 0.6696 | 0.6723 |
| | | R-square | 0.2128 | 0.3180 | 0.3205 | 0.3175 |
| | | MSE for beta | 0.2328 | 0.2209 | 0.2190 | 0.2244 |
| | | Main Effect TPR | 0.524 | 0.605 | 0.606 | 0.609 |
| | | Main Effect FPR | 0.0950 | 0.1220 | 0.1224 | 0.1296 |
| | | Interaction TPR | 0.1470 | 0.5258 | 0.8021 | 1.0624 |

Table S5: *Results of simulations in Section 3: summarized from 100 simulations under setting (a) and (b) with* **n=2000, p=100.**

| **Setting** | | | **r=2** | **r=5** | **r=8** | **r=10** |
|---|---|---|---|---|---|---|
| Setting (a)<br>$\mathbf{x}_i = \Lambda\mathbf{z}_i + \mathbf{u}_i$<br>$\Lambda = [\mathbf{l_1}, \ldots, \mathbf{l_1}]$<br>$\mathbf{l}_1 = \text{diag}(\sqrt{0.8})_{K\times K}$ | LF-VCR | MSE | 0.7092 | 0.6199 | 0.5651 | 0.5452 |
| | | R-square | 0.2917 | 0.3808 | 0.4357 | 0.4556 |
| | | MSE for beta | 0.2466 | 0.2155 | 0.1785 | 0.1558 |
| | | Main Effect TPR | 0.988 | 0.992 | 0.994 | 0.997 |
| | | Main Effect FPR | 0.2141 | 0.2258 | 0.2343 | 0.2400 |
| | | Interaction TPR | 0.2827 | 0.3863 | 0.3906 | 0.3882 |
| | | Interaction FPR | 0.2697 | 1.4924 | 2.6370 | 3.3734 |
| | LF-VCR (ae) | MSE | 0.7191 | 0.6189 | 0.5638 | 0.5465 |
| | | R-square | 0.2818 | 0.3823 | 0.4372 | 0.4545 |
| | | MSE for beta | 0.2513 | 0.2191 | 0.1784 | 0.1563 |
| | | Main Effect TPR | 0.982 | 0.990 | 0.993 | 0.995 |
| | | Main Effect FPR | 0.2220 | 0.2314 | 0.2356 | 0.2397 |
| | | Interaction TPR | 0.2801 | 0.3967 | 0.4038 | 0.3902 |
| | | Interaction FPR | 0.2719 | 1.5360 | 2.7332 | 3.3956 |
| Setting (b)<br>$\mathbf{x}_i = \Lambda\mathbf{z}_i + \mathbf{u}_i$<br>$\Lambda = \min_\Lambda \left\lVert \Lambda\Lambda^T - A \right\rVert_F$<br>$A = Q\Lambda_{\text{diag}}Q^T$ | LF-VCR | MSE | 0.7620 | 0.6503 | 0.6547 | 0.6578 |
| | | R-square | 0.2417 | 0.3524 | 0.3485 | 0.3457 |
| | | MSE for beta | 0.2087 | 0.1781 | 0.1779 | 0.1789 |
| | | Main Effect TPR | 0.745 | 0.826 | 0.812 | 0.801 |
| | | Main Effect FPR | 0.1280 | 0.1360 | 0.1273 | 0.1230 |
| | | Interaction TPR | 0.2109 | 0.7142 | 1.0973 | 1.3375 |
| | LF-VCR (ae) | MSE | 0.7628 | 0.6521 | 0.6518 | 0.6532 |
| | | R-square | 0.2413 | 0.3507 | 0.3512 | 0.3498 |
| | | MSE for beta | 0.2075 | 0.1826 | 0.1810 | 0.1846 |
| | | Main Effect TPR | 0.742 | 0.828 | 0.829 | 0.832 |
| | | Main Effect FPR | 0.1260 | 0.1460 | 0.1423 | 0.1427 |
| | | Interaction TPR | 0.2013 | 0.6749 | 0.9841 | 1.2595 |

Table S6: *Results of simulations in Section 3: summarized from 100 simulations under setting (a) and (b) with* **n=4000, p=100.**

| **Setting** | | | **r=2** | **r=5** | **r=8** | **r=10** |
|---|---|---|---|---|---|---|
| Setting (a)<br>$\mathbf{x}_i = \Lambda \mathbf{z}_i + \mathbf{u}_i$<br>$\Lambda = [\mathbf{l_1}, \ldots, \mathbf{l_1}]$<br>$\mathbf{l}_1 = \text{diag}(\sqrt{0.8})_{K\times K}$ | LF-VCR | MSE | 0.6852 | 0.5873 | 0.5303 | 0.5121 |
| | | R-square | 0.3171 | 0.4142 | 0.4719 | 0.4905 |
| | | MSE for beta | 0.1970 | 0.1585 | 0.1149 | 0.0892 |
| | | Main Effect TPR | 1.000 | 1.000 | 1.000 | 1.000 |
| | | Main Effect FPR | 0.2240 | 0.2398 | 0.2321 | 0.2341 |
| | | Interaction TPR | 0.3475 | 0.4412 | 0.4160 | 0.3830 |
| | | Interaction FPR | 0.3377 | 1.7109 | 2.8227 | 3.3314 |
| | LF-VCR (ae) | MSE | 0.6863 | 0.5909 | 0.5292 | 0.5126 |
| | | R-square | 0.3161 | 0.4111 | 0.4732 | 0.4898 |
| | | MSE for beta | 0.2005 | 0.1611 | 0.1174 | 0.0897 |
| | | Main Effect TPR | 1.000 | 1.000 | 1.000 | 1.000 |
| | | Main Effect FPR | 0.2280 | 0.2340 | 0.2345 | 0.2360 |
| | | Interaction TPR | 0.3604 | 0.4351 | 0.4260 | 0.3897 |
| | | Interaction FPR | 0.3481 | 1.6893 | 2.8881 | 3.3916 |
| Setting (b)<br>$\mathbf{x}_i = \Lambda \mathbf{z}_i + \mathbf{u}_i$<br>$\Lambda = \min_\Lambda \left\|\Lambda\Lambda^T - A\right\|_F$<br>$A = Q\Lambda_{\text{diag}}Q^T$ | LF-VCR | MSE | 0.7504 | 0.6359 | 0.6382 | 0.6395 |
| | | R-square | 0.2536 | 0.3674 | 0.3653 | 0.3641 |
| | | MSE for beta | 0.1829 | 0.1470 | 0.1498 | 0.1503 |
| | | Main Effect TPR | 0.932 | 0.961 | 0.954 | 0.948 |
| | | Main Effect FPR | 0.1607 | 0.1770 | 0.1647 | 0.1528 |
| | | Interaction TPR | 0.2931 | 1.0264 | 1.5216 | 1.7543 |
| | LF-VCR (ae) | MSE | 0.7504 | 0.6370 | 0.6371 | 0.6383 |
| | | R-square | 0.2535 | 0.3665 | 0.3662 | 0.3651 |
| | | MSE for beta | 0.1843 | 0.1527 | 0.1518 | 0.1557 |
| | | Main Effect TPR | 0.934 | 0.961 | 0.961 | 0.959 |
| | | Main Effect FPR | 0.1749 | 0.1900 | 0.1888 | 0.1867 |
| | | Interaction TPR | 0.3178 | 0.9589 | 1.4845 | 1.8687 |

Table S7: *Results of simulations in Section 3: summarized from 100 simulations under setting (a) and (b) with* **n=6000, p=100.**

| **Setting** | | | **r=2** | **r=5** | **r=8** | **r=10** |
|---|---|---|---|---|---|---|
| Setting (a)<br>$\mathbf{x}_i = \Lambda \mathbf{z}_i + \mathbf{u}_i$<br>$\Lambda = [\mathbf{l_1}, \ldots, \mathbf{l_1}]$<br>$\mathbf{l}_1 = \text{diag}(\sqrt{0.8})_{K\times K}$ | LF-VCR | MSE | 0.6825 | 0.5775 | 0.5203 | 0.5019 |
| | | R-square | 0.3183 | 0.4233 | 0.4809 | 0.4997 |
| | | MSE for beta | 0.1797 | 0.1343 | 0.0927 | 0.0636 |
| | | Main Effect TPR | 1.000 | 1.000 | 1.000 | 1.000 |
| | | Main Effect FPR | 0.2364 | 0.2391 | 0.2360 | 0.2295 |
| | | Interaction TPR | 0.3804 | 0.4613 | 0.4325 | 0.3773 |
| | | Interaction FPR | 0.3684 | 1.7873 | 2.9378 | 3.2829 |
| | LF-VCR (ae) | MSE | 0.6784 | 0.5765 | 0.5189 | 0.5021 |
| | | R-square | 0.3221 | 0.4243 | 0.4822 | 0.4994 |
| | | MSE for beta | 0.1818 | 0.1385 | 0.0931 | 0.0639 |
| | | Main Effect TPR | 1.000 | 1.000 | 1.000 | 1.000 |
| | | Main Effect FPR | 0.2324 | 0.2376 | 0.2384 | 0.2325 |
| | | Interaction TPR | 0.3896 | 0.4726 | 0.4326 | 0.3847 |
| | | Interaction FPR | 0.3770 | 1.8323 | 2.9290 | 3.3533 |
| Setting (b)<br>$\mathbf{x}_i = \Lambda \mathbf{z}_i + \mathbf{u}_i$<br>$\Lambda = \min_\Lambda \left\|\Lambda\Lambda^T - A\right\|_F$<br>$A = Q\Lambda_{\text{diag}}Q^T$ | LF-VCR | MSE | 0.7475 | 0.6295 | 0.6308 | 0.6317 |
| | | R-square | 0.2568 | 0.3741 | 0.3731 | 0.3722 |
| | | MSE for beta | 0.1689 | 0.1296 | 0.1314 | 0.1328 |
| | | Main Effect TPR | 0.981 | 0.993 | 0.992 | 0.988 |
| | | Main Effect FPR | 0.1676 | 0.2107 | 0.1780 | 0.1754 |
| | | Interaction TPR | 0.3348 | 1.3459 | 1.7534 | 2.1669 |
| | LF-VCR (ae) | MSE | 0.7477 | 0.6306 | 0.6307 | 0.6316 |
| | | R-square | 0.2566 | 0.3730 | 0.3729 | 0.3721 |
| | | MSE for beta | 0.1694 | 0.1365 | 0.1346 | 0.1394 |
| | | Main Effect TPR | 0.981 | 0.992 | 0.995 | 0.993 |
| | | Main Effect FPR | 0.1764 | 0.2261 | 0.2148 | 0.2164 |
| | | Interaction TPR | 0.3505 | 1.2715 | 1.8365 | 2.2351 |
